## Supplementary Figures for "Age Influences on the Molecular Presentation of Tumours"

**Supplementary Figures 1-5**

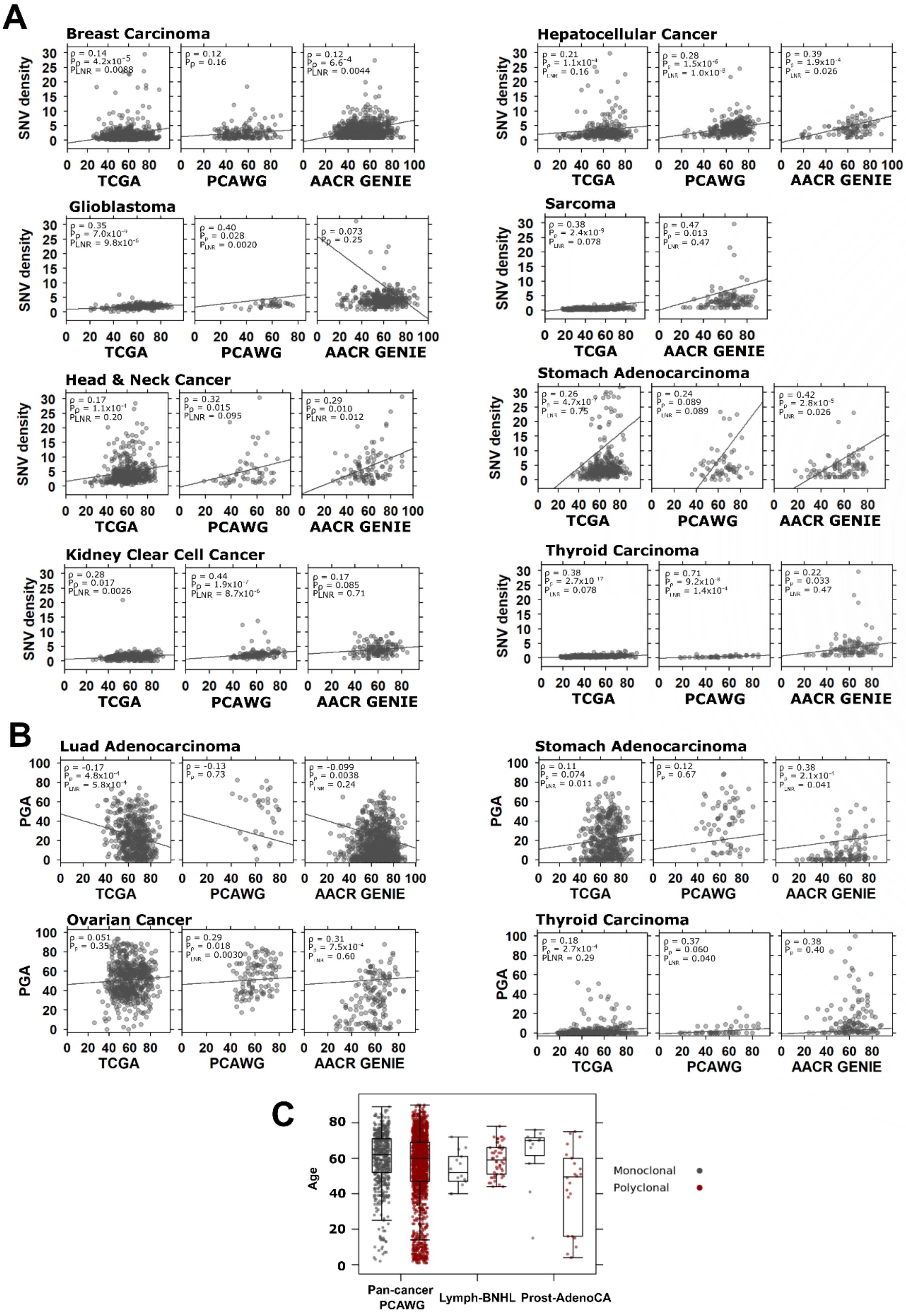

**Supplementary Figure 1. Age-associated differences in PGA and SNV density.**

Associations of age and **(A)** SNV density and **(B)** PGA compared across three datasets (when available). Spearman correlation coefficient (ρ), adjusted Spearman and multivariate linear regression p-values shown. **(C)** Age compared between monoclonal and polyclonal tumours in pan-cancer PCAWG, non-Hodgkin Lymphoma, and prostate cancer. Tukey boxplots are shown with the box indicating quartiles and the whiskers drawn at the lowest and highest points within 1.5 interquartile range of the lower and upper quartiles, respectively.

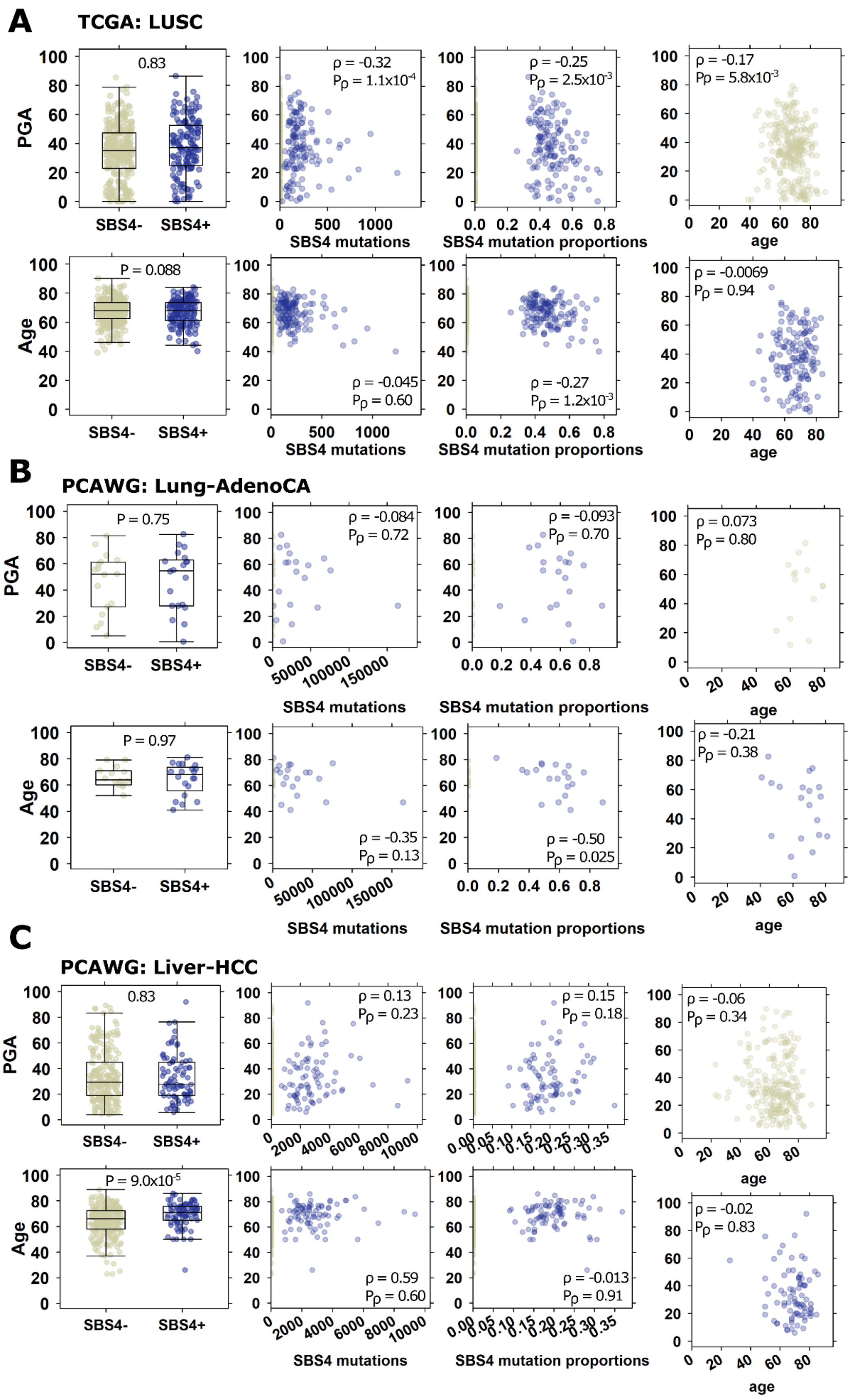
A
TCGA: LUSC
PGA
Age
SBS4-
SBS4+
0.83
ρ = -0.32
Pρ = 1.1x10-4
ρ = -0.25
Pρ = 2.5x10-3
ρ = -0.17
Pρ = 5.8x10-3
P = 0.088
ρ = -0.045
Pρ = 0.60
ρ = -0.27
Pρ = 1.2x10-3
ρ = -0.0069
Pρ = 0.94
SBS4 mutations
SBS4 mutation proportions
age
B
PCAWG: Lung-AdenoCA
P = 0.75
ρ = -0.084
Pρ = 0.72
ρ = -0.093
Pρ = 0.70
ρ = 0.073
Pρ = 0.80
P = 0.97
ρ = -0.35
Pρ = 0.13
ρ = -0.50
Pρ = 0.025
ρ = -0.21
Pρ = 0.38
C
PCAWG: Liver-HCC
0.83
ρ = 0.13
Pρ = 0.23
ρ = 0.15
Pρ = 0.18
ρ = -0.06
Pρ = 0.34
P = 9.0x10-5
ρ = 0.59
Pρ = 0.60
ρ = -0.013
Pρ = 0.91
ρ = -0.02
Pρ = 0.83

**Supplementary Figure 2. Associations of age and PGA by mutational signature SBS4.**

Associations of age and PGA with tobacco-related signature SBS4 vary by tumour-type as shown for **(A)** TCGA lung squamous cell carcinoma, **(B)** PCAWG lung adenocarcinoma, and **(C)** PCAWG hepatocellular carcinoma. From left to right, plot columns show comparisons of PGA and Age with SBS4 detection, number of SBS-4 attributed mutations, relative proportion of SBS4-attributed mutations, and PGA *vs*. age broken down by SBS4-status.

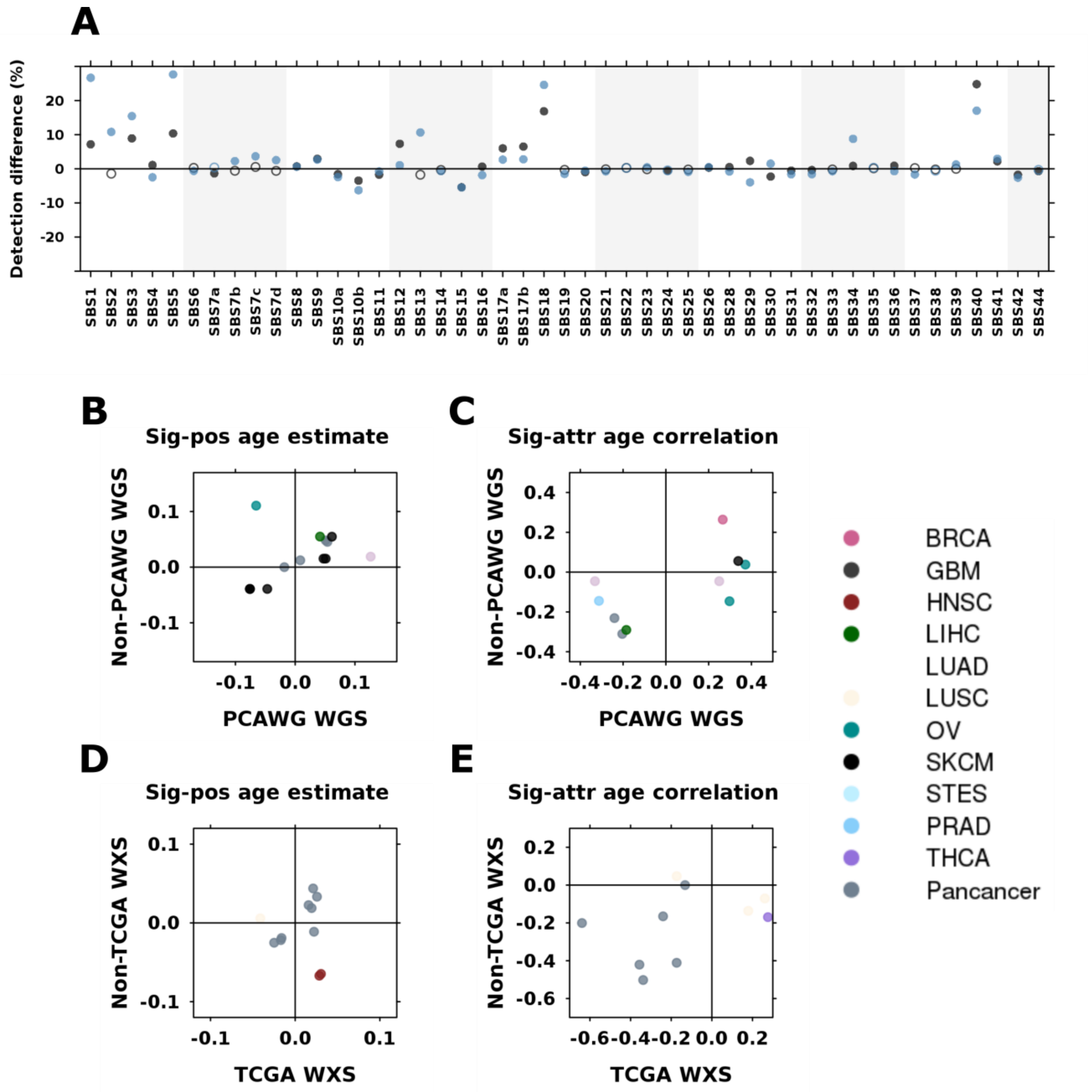


**Supplementary Figure 3. Validation of age-biased signatures in external datasets.**

**(A)** For each signature, we compared the detection difference between whole genome and exome sequencing. Black dots show the difference in detection between PCAWG WGS and TCGA WXS. Blue dots show the difference in detection between non-PCAWG WGS and non-TCGA WXS. Closed marks indicate comparisons where the difference is statistically significant (proportion test $p < 0.05$) and open marks indicate non-significant comparisons. We use non-PCAWG whole genome sequencing and non-TCGA whole exome sequencing data to examine the effect sizes and direction of signatures found to be age-biased in (**B-C**) PCAWG and (**D-E**) TCGA data. **(B)** and **(D)** show the logistic regression coefficient estimates for signatures with age-biased detection. **(C)** and **(E)** show Spearman's correlation between signature relative activity and age.

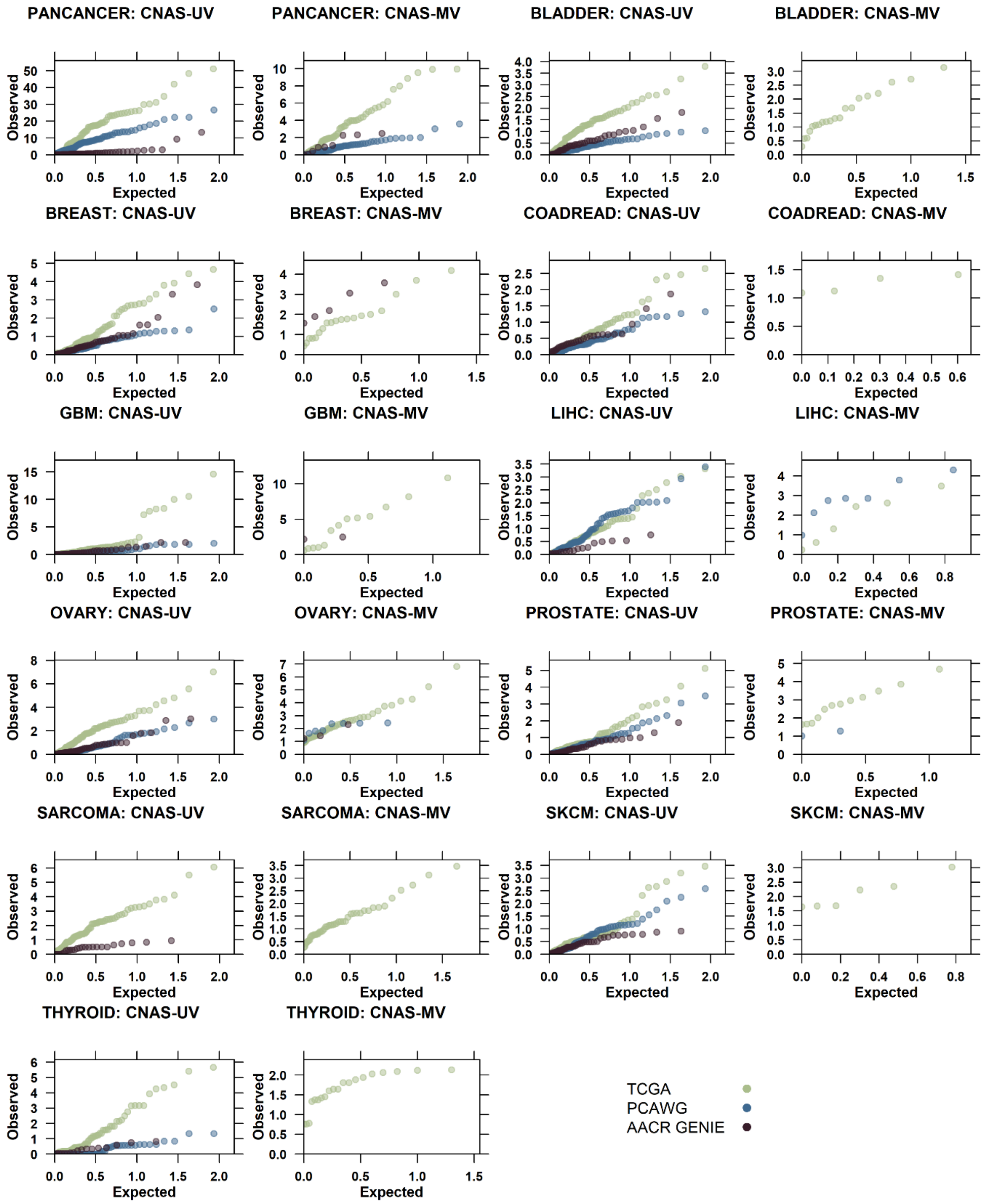


**Supplementary Figure 4. Q-Q plots of CNA-age p-values *vs*. expected uniform distribution.**

Comparison of -$\log_{10}$(p-value) distributions for univariate (UV) and multivariate (MV) CNA-age association tests against expected univariate distributions. Only tumour-types with significant multivariate results shown (as seen in **Figure 3A-C**).

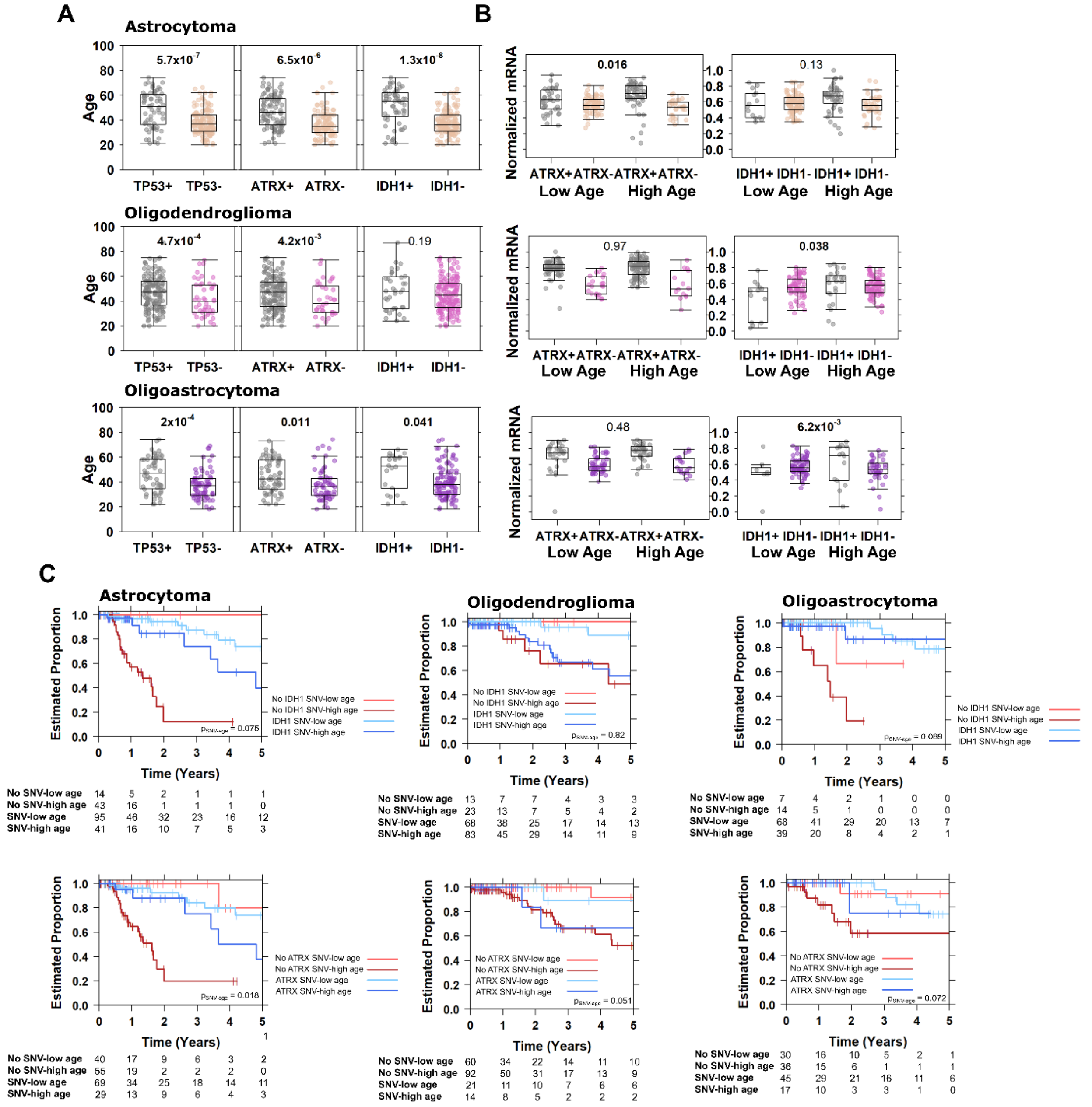


**Supplementary Figure 5. Age-associated driver differences by lower grade glioma subtype.**

**(A)** Age-associated drivers identified in lower grade glioma have subtype-dependent relationships. Associations of *ATRX* and *IDH1* mutation with **(B)** mRNA and **(C)** overall survival are also dependent on subtype.
